# Multiple lineages of transmissible neoplasia in the basket cockle (*C. nuttallii*) with repeated horizontal transfer of mitochondrial DNA

**DOI:** 10.1101/2023.10.11.561945

**Authors:** Marisa A. Yonemitsu, Jordana K. Sevigny, Lauren E. Vandepas, James L. Dimond, Rachael M. Giersch, Helen J. Gurney-Smith, Cathryn L. Abbott, Janine Supernault, Ruth Withler, Peter D. Smith, Sydney A. Weinandt, Fiona E. S. Garrett, Zachary J. Child, Robin Little Wing Sigo, Elizabeth Unsell, Ryan N. Crim, Michael J. Metzger

**Author notes:** these authors contributed equally (Marisa A. Yonemitsu, Jordana K. Sevigny). Corresponding author: Michael J. Metzger.

## Abstract

Transmissible cancers are clonal lineages of neoplastic cells able to infect multiple hosts, spreading through populations in the environment as an infectious disease. Transmissible cancers have been identified in Tasmanian devils, dogs, and bivalves. Several lineages of bivalve transmissible neoplasias (BTN) have been identified in multiple bivalve species. In 2019 in Puget Sound, Washington, USA, disseminated neoplasia was observed in basket cockles (*Clinocardium nuttallii*), a species that is important to the culture and diet of the Suquamish Tribe as well as other tribes with traditional access to the species. To test whether disseminated neoplasia in cockles is a previously unknown lineage of BTN, a nuclear locus was amplified from cockles from Agate Pass, Washington, and sequences revealed evidence of transmissible cancer in several individuals. We used a combination of cytology and quantitative PCR to screen collections of cockles from eleven locations in Puget Sound and along the Washington coastline to identify the extent of contagious cancer spread in this species. Two BTN lineages were identified in these cockles, with one of those lineages (CnuBTN1) being the most prevalent and geographically widespread. Within the CnuBTN1 lineage, multiple nuclear loci support the conclusion that all cancer samples form a single clonal lineage. However, the mitochondrial alleles in each cockle with CnuBTN1 are different from each other, suggesting mitochondrial genomes of this cancer have been replaced multiple times during its evolution, through horizontal transmission. The identification and analysis of these BTNs are critical for broodstock selection, management practices, and repopulation of declining cockle populations, which will enable continued cultural connection and dietary use of the cockles by Coast Salish Tribes.

## INTRODUCTION

Cancer arises from oncogenic mutations within the cells of an organism and, despite being able to grow and metastasize, most cancers are confined to live and die within the body in which they originate. However, there are rare cases of naturally occurring transmissible cancers that have evolved to spread between individuals as an infectious allograft, causing disease long after their original host is gone (Metzger & Goff, 2016). This phenomenon has been observed once in dogs (Murgia et al., 2006; Rebbeck et al., 2009), twice in Tasmanian devils (Pearse & Swift, 2006; Pye et al., 2016), and at least eight times in several marine bivalve species (Garcia-Souto et al., 2022; Metzger et al., 2015; Metzger et al., 2016; Michnowska et al., 2022; Yonemitsu et al., 2019).

In bivalves, disseminated neoplasia (DN) is a leukemia-like disease characterized by a massive proliferation of rounded, polyploid cells in the bivalve hemolymph (circulatory fluid) that eventually progresses to cancer dissemination into host tissues and usually results in death of the individual. Outbreaks of DN in marine bivalves across the globe have historically led to massive die-offs of affected populations, posing a threat to commercial aquaculture and species conservation efforts (Barber, 2004; Carballal et al., 2015; Farley et al., 1991; Muttray et al., 2012). Of the bivalve species affected by outbreaks of DN, recent genetic analyses of DN in soft-shell clams (*Mya arenaria* (Metzger et al., 2015)), mussels (*Mytilus trossulus, M. edulis, M. chilensis, M. galloprovincialis* (Hammel et al., 2021; Metzger et al., 2016; Yonemitsu et al., 2019)), common cockles (*Cerastoderma edule* (Metzger et al., 2016)), golden carpet shell clams (*Polititapes aureus* (Metzger et al., 2016)), baltic clams (*Macoma balthica*(Michnowska et al., 2022)), and warty venus clams (*Venus verrucosa* (Garcia-Souto et al., 2022)) have shown that the DN affecting these species are cases of bivalve transmissible neoplasia (BTN), suggesting that transmissible cancer may be a more common phenomenon in marine invertebrates than previously recognized.

The basket cockle, *Clinocardium nuttallii*, is a culturally significant shellfish species for the Suquamish Tribe, as well as many other coastal tribes from Washington to Alaska, who engage in subsistence foraging of native shellfish species as a cultural and dietary practice (Poe et al., 2016). *C. nuttallii* occurs in both intertidal and subtidal habitats with a wide range along the Pacific coast from southern California to western Alaska and Japan (Ahyong et al., 2023; Barber et al., 2019). A long-term survey of Washington *C. nuttallii* populations showed varied population dynamics across beaches in the southern Salish Sea with decreased population abundance observed on some beaches, while other locations maintained steady numbers or increased in population size (Barber et al., 2019). However, Suquamish Tribal members have noted declines in local subsistence basket cockle populations over the last two decades. The cultural importance of this species has driven efforts to investigate *C. nuttallii* aquaculture production programs to supplement wild populations.

During health assessments of potential *C. nuttallii* broodstock populations, several individuals were diagnosed with DN through histological analysis (Ralph Elston, AquaTechnics, personal communication). This suggested that DN may have an impact on the health and viability of *C. nuttallii* aquaculture strategies, given the negative impact of DN on commercial shellfish farms, and raised the question of whether the DN in basket cockles is a transmissible cancer (BTN).

While moving toward hatchery production of cockles, the Suquamish Tribe and the Puget Sound Restoration Fund simultaneously pursued research looking into whether genetic population structure exists in cockle populations of Washington State. Collections from eleven locations were performed throughout Washington State with permission and assistance from multiple Tribes (Dimond et al., 2022). The cockles from these collections were also sampled for the presence of DN.

In this study, we investigated the DN found in *C. nuttallii* to determine whether the cancer is a BTN. Combining the analysis of multiple nuclear genetic markers with cytology of cells in the cockle hemolymph, we identified two independent lineages of *C. nuttallii* -derived BTN, which we term CnuBTN1 and 2. We developed qPCR-based assays to screen for both cancer lineages in cockle populations collected from beaches across the Salish Sea and Washington’s Pacific coast, finding this cancer in cockle populations at three of eleven locations. Interestingly, analysis of multiple mitochondrial DNA (mtDNA) markers showed extensive mitochondrial replacement occurring in the most widespread lineage, CnuBTN1, an observation that has been made in three other transmissible cancers, including the canine transmissible venereal tumor (CTVT) in dogs (Rebbeck et al., 2011; Strakova et al., 2016; Strakova et al., 2020) and BTNs in *M. chilensis* (Yonemitsu et al., 2019) and *C. edule* (Bruzos et al., 2023), suggesting that altered mitochondrial dynamics may be an important factor in the evolution of these cancers.

## MATERIALS AND METHODS

### Sample collection

Pilot study samples were collected in March 2019 at low tide from the intertidal zone of Agate Pass, WA with a shovel or clam rake (Table S1). Cockles were anesthetized in 50g/L MgCl × 6H_2_O in artificial seawater (Instant Ocean) for 1 hr until the valves opened and did not close when pressure was applied. One half to 1 mL of hemolymph was extracted from the anterior or posterior adductor muscle of each animal using a 1 inch 26-gauge needle. Several drops of hemolymph were placed in a well of a 96-well plate and incubated for about 1 hr at 4°C to allow normal cells to adhere to the plate. Samples were then screened visually for obvious disseminated neoplasia using a phase contrast inverted microscope, as previously described (Giersch et al., 2022). The remaining hemolymph was spun down at 1,000×g for 10 minutes at 4°C to pellet the cells and remove cell-free hemolymph. Animals were then sacrificed and ∼50 mg of mantle tissue was collected and frozen at -80°C either in ethanol or in RNAlater (Invitrogen) until DNA extraction.

Additional cockles from 11 intertidal collections from across the Salish Sea and Pacific coast of Washington were sampled in October 2019 to January 2020 (Table S1), as described previously (Dimond et al., 2022). For hemolymph collection from animals without MgCl treatment, the syringe was placed between the edges of the valves, and, if needed, a small hole was bored into the anterior edge of the valves and 0.5-1 mL of hemolymph was collected from the muscle and processed as above.

### Cytology

Fixing of cells for histology was done with fresh hemolymph, as described in detail previously. Briefly, 100 µL of the collected hemolymph was applied to poly-L-lysine slides (Electron Microscopy Sciences, Hatfield, PA, USA) and allowed to rest for ten minutes at room temperature on a flat surface. A “blood smear” was performed by gently removing excess hemolymph using a coverslip and air drying. The adhered hemocytes were then fixed in 100% methanol for 15 minutes. The slides were removed from methanol, air dried at room temperature, and stained with hematoxylin & eosin (Leica, USA). Cytology was conducted for all cockles, excluding the March 2019 cockles from Agate Pass (Y), and the cockles from Hood Canal (HC), Eld Inlet (SS), and cockles 13-30 from Squaxin Island (SX).

### DNA extraction

DNA was extracted from ethanol-fixed mantle and hemocyte samples using DNeasy Blood & Tissue Kit (Qiagen) (Dimond et al., 2022). Mantle extractions of the pilot samples were performed with additional steps to remove PCR-inhibiting polysaccharides: After tissue lysis, 65 µl of P3 Buffer (Qiagen) was added for 5 minutes to precipitate out polysaccharides and spun down at 17,000×g at 4°C for 5 minutes. The resulting supernatant was transferred to a new tube, 200 µl of Buffer AL was added, and the manufacturer’s protocol resumed.

### α-tubulin, mtCOI, and mtCYTB PCR and cloning

Primers and annealing temperatures are listed in Table S2. Without a reference genome available, we searched for primers that would consistently amplify a variable nuclear locus by targeting conserved genes with available sequence from a related bivalve species (usually *Cerastoderma edule*), and we tested multiple primers for each gene that were designed to target intron-spanning coding regions. After testing multiple candidate nuclear loci (*α-amylose*, *RNA Pol II*, *α-tubulin, β-actin, H3, ITS-1, and 16S*), only one (*α-tubulin*) met our criteria: the correct locus amplified in all samples, 1-2 alleles were identified in all healthy samples, and polymorphisms were detectable in the healthy population. PCR amplification at the *α-tubulin* locus was done using Q5 Hot Start High-Fidelity DNA Polymerase (NEB) with a 30 s extension time. Primers for *α-tubulin* were designed in exons flanking an intron to amplify a ∼850 bp intron-spanning region. Primers for *cytochrome B* were designed to amplify a ∼500 bp region. About 25 ng of genomic DNA was amplified for 35 cycles. PCR products were gel extracted using spin columns (Qiagen) and sequenced, or, when multiple alleles at a locus could not be resolved by direct sequencing, were cloned using the Zero Blunt TOPO PCR Cloning Kit (Invitrogen). Plasmids were transformed into DH5α competent *E. coli* (Invitrogen) and 6-12 clones were picked for further sequencing (Azenta/Genewiz). The primer binding regions were excluded from sequence analysis. Allele-specific SNVs were identified after cloning of *α-tubulin* alleles. PCR products for all *α-tubulin* amplicons and for the *mtCYTB* samples from Y504 were also sequenced using Nanopore rapid barcoding chemistry, and after assembly of the consensus sequence, the number of reads supporting each SNV were determined by aligning the raw reads to the consensus (sequencing and mapping done by Primordium Labs/Plasmidsaurus, USA). For alleles with multiple unique SNVs, the frequencies of each SNV corresponding to a particular allele were averaged to determine the allele frequency. One allele lacked any unique SNVs (e), but it was different from the k allele by only a single SNV; in this case, the e allele frequency for each sample was calculated by taking the average frequency of those SNVs that were shared between k and e in those samples and subtracting the frequency of the SNV found only in k and not e (Figure S1).

### Phylogenetic trees

All sequences were manually aligned in BioEdit (Hall, T.A., 1999). Maximum likelihood phylogenies were built by PhyML 3.0 (Guindon et al., 2010) using Akaike Information Criterion to select the most suitable substitution model (Lefort et al., 2017) with a standard bootstrap analysis (100 repeats). The phylogenetic trees were visualized in FigTree v1.4.4 (http://tree.bio.ed.ac.uk/software/figtree/).

*α-tubulin* allele-specific qPCR:

Cancer-associated, allele-specific qPCR assays were developed and performed in *C. nuttallii* genomic DNA samples using PowerUp SYBR Green Master Mix (Applied Biosystems). Primer sets were designed to specifically amplify each of the cancer-associated *α-tubulin* alleles, allele k (CnuBTN1) and allele p (CnuBTN2). A pair of universal control primers were designed to amplify all possible *α-tubulin* alleles. Primers are listed in Table S3. Standards for all targets consisted of a single plasmid containing the relevant, cancer-associated allele target, which was linearized by NotI-HF (NEB) restriction digest before qPCR. Serial dilutions (10^7^-10^1^ copies) of linearized standard plasmid in triplicate were used to generate a standard curve for each qPCR run (quantified by qubit). In all experimental cases, 5-50 ng of each genomic DNA sample was run in triplicate.

Allele-specific qPCR using *α-tubulin* k and *α-tubulin* universal control primers used a 3-step cycling stage (95°C for 15s, 50°C for 15s, and 60°C for 30s, respectively) for 40 cycles, whereas qPCR using *α-tubulin* p primers was run using a 2-step cycling stage protocol (95°C for 15s, 60°C for 30s). The ratio of the cancer-associated allele to all alleles present at a given locus was calculated for each sample using the average copy number of cancer-associated allele divided by the average copy number of total alleles to generate an estimate of the fraction of total alleles that are the cancer-associated allele. Animals were identified as potentially diseased if the ratio of hemocyte and tissue cancer-associated allele fractions was at least 1.5 in CnuBTN1 (k), due to the finding of animals with the k allele in the host genome, and at least 0.5 for CnuBTN2 (p). If the control qPCR amplified below a threshold of 1,000 copies per reaction, the sample was excluded due to low DNA extraction yield.

### Identification of microsatellite primers

Following a microsatellite identification method used previously (Abbott et al., 2011), in 2010, genomic libraries were constructed from adductor muscle DNA from one individual cockle (DNeasy Tissue Kit, Qiagen), from Deep Bay, British Columbia, Canada (49°27’35”N, 124°44’02”W), eluted with molecular grade water. The resulting DNA was quality checked and submitted to the McGill University and Genome Quebec Innovation Centre for 454 sequencing using a Roche GS-FLX sequencer. Approximately 12% of 454 reads were assembled into 2,597 contigs with mean length of 643 bp. A custom Perl script was used to search contigs and raw reads for uninterrupted microsatellites with repeat motifs 2-10 bases long; duplicate loci were excluded. PCR primers were designed using Primer3 (Rozen and Skaletsky 2000) for 89 unique microsatellites to test for polymorphism that had <20 bp of flanking sequence and seven or more repeat units in 44 contigs. We also found 396 di- or tri-microsatellites of 14 or more repeats and with <30 bp of flanking sequence in 374 unassembled 454 reads. Sequence reads were also mined for tetranucleotide repeats with at least 14 repeat units and 30 bp of flanking sequence; we found 68 of these, each in a unique read. Sequences used for primer design were aligned and duplicates were excluded (Sequencher v 4.8, Gene Codes Corp.). From this, PCR primers were designed and used to screen 89 unique microsatellites.

For the initial testing of microsatellite primers, PCR amplifications used a DNA Engine Tetrad2 thermal cycler (BioRad, Hercules, California) in 5 µL volumes consisting of 0.14 U of Qiagen HotStar Taq DNA polymerase, 1 µL 1:8 diluted DNA, 1× PCR buffer, 84 µM each dNTP, 0.50 µM M13 fluorescently-labeled primer, 0.13 µM M13 labeled forward primer, 0.50 µM reverse primer, and deionized H_2_O. The thermal cycling profile included 15 min at 95°C followed by 8 cycles of 30 s at 94°C, 30s at 51°C and 30-40s at 72°C. Microsatellites were run on an ABI 3730, and genotypes scored with ABI GeneMapper version 4.0 using an internal lane sizing standard (GS500LIZ_3730, Applied Biosystems). A temperature gradient with annealing temperatures ranging from 48-59°C increasing in 1°C increments was used to determine optimal annealing temperature for each locus. Initially, all 89 loci were tested for polymorphism on eight cockle samples from three sample sites in Canada, after which a subset of 20 loci were selected for additional testing on 300 basket cockles from three locations. Based on the level of polymorphism, allele size range, success of amplification, and ease of scoring, 8 loci were chosen for further use. Microsatellite details, primer sequences and optimal annealing temperatures for these loci are given in Table S4.

### Analysis of microsatellite loci

Microsatellites were amplified from DNA extracted from both hemocyte and tissue using Taq polymerase (Genesee Scientific) for a select number of individuals: all possible neoplastic animals positive by the qPCR screening criteria (n=22) and one randomly selected, non-neoplastic animal from each collection (n=11), and all six animals from the pilot collection (n=3 healthy, n=3 with BTN). Microsatellites were amplified using primers for 8 polymorphic loci: Cnu48, Cnu55, Cnu58, Cnu63, Cnu68, Cnu72, Cnu78, and Cnu81 (Table S4). Allele sizes were identified by fragment analysis using fluorescent primers (6-FAM, PET, NED, and VIC) and a 3730xl Genetic Analyzer with the LIZ-500 size standard (Applied Biosystems, operated by Genewiz). Alleles were called using Peak Scanner 2.0 (Applied Biosystems; see (Metzger et al., 2016)), and rounded to the nearest integer and binned using TANDEM (Matschiner & Salzburger, 2009) to account for split rounding of decimals and assay variability (Figure S2 and Table S4). During integer correction, one population of origin was assumed with ploidy set to two, though for polyploid animals, the samples were split into two rows as if they were different samples to accommodate all the alleles.

After integer correction, the microsatellites were analyzed using R package, *PoppR,* Version 2.9.6 (Kamvar et al., 2014), employing Bruvo’s method with the infinite alleles model to build distance matrices for phylogenetic tree creation. The variable “c” (repeat size) was set to 0.00001 resulting in the distance between two alleles to be 0 (identical) or 1 (non-identical), adhering to the band sharing model (Metzger et al., 2015). The variable “sample” (number of bootstrap replicates) was set to 1000. All samples were coded to be from one population. We used two methods to calculate genetic distance. For generation of a phylogenetic tree, we used a binary presence/absence model. Each allele was classified as a dominant marker with a total of 136 markers in 8 loci (Metzger et al., 2015; Rodzen & May, 2002). Each peak (allele) was coded as present (1) or absent (2). In normal individuals, identical peaks were observed in the hemocyte DNA and tissue DNA and they were therefore considered present in both. In highly neoplastic animals (those with more cancer cells than host cells in the hemolymph), the cancer-associated alleles were present in the hemocyte DNA and to a lesser extent in the host solid tissue; occasionally the cancer-associated alleles obscured possible host genotypes, resulting in a potentially ambiguous call. We conducted an alternative sub-analysis in which any peak that might be ambiguous was coded as missing (0) and excluded from pairwise calculations. The tree that resulted from this alternate binary analysis had the same topology of the tree using the most likely genotype, showing that these calls do not affect the conclusions. The distance between two samples in this distance matrix is the number of differences in the pairwise comparison at each observed allele divided by the number of all observed alleles, excluding comparisons in which one or both individuals have an ambiguous value. The phylogenetic tree generated by *PoppR* was visualized using FigTree v1.4.4.

For direct measurement of pairwise distances, we used a co-dominant polyploid model. In this model, each locus is coded with all the allele sizes observed. In normal individuals, one or two sizes were observed for each locus, but in the cancer samples up to four alleles were observed. Therefore, the ploidy was set to four, and any samples with fewer alleles had “0” indicating missing data for the remaining alleles at that locus. These missing values are ignored in the analysis using Bruvo’s method, which was designed to compare samples of different ploidy levels. This generates a distance that is proportional to the number of loci, rather than the number of observed alleles, and is a more accurate direct measure of genetic distance.

## RESULTS

### Disseminated neoplasia in cockles is a bivalve transmissible neoplasia

Due to the recent identification of DN in Puget Sound basket cockles (*C. nuttallii*), found through histological examination during health screening of potential broodstock, we set out to determine whether the DN was a transmissible cancer, as seen in other bivalves. We first examined hemolymph samples from cockles collected from Agate Pass, Washington (Figure 1A), a site where DN had previously been observed in cockles through histological analysis (Vandepas et al., 2024). Hemocytes of non-neoplastic cockles are highly adherent to tissue culture dishes and show a variety of cell types, many with pseudopodia (Figure 1B). In this collection of cockles from Agate Pass, we identified three individuals with DN, in which nearly all cells in the hemolymph display the rounded, uniform, non-adherent morphology typical of DN cells (Barber, 2004; Carballal et al., 2015; Metzger et al., 2015; Metzger et al., 2016) (Figure 1C).

**Figure 1.**
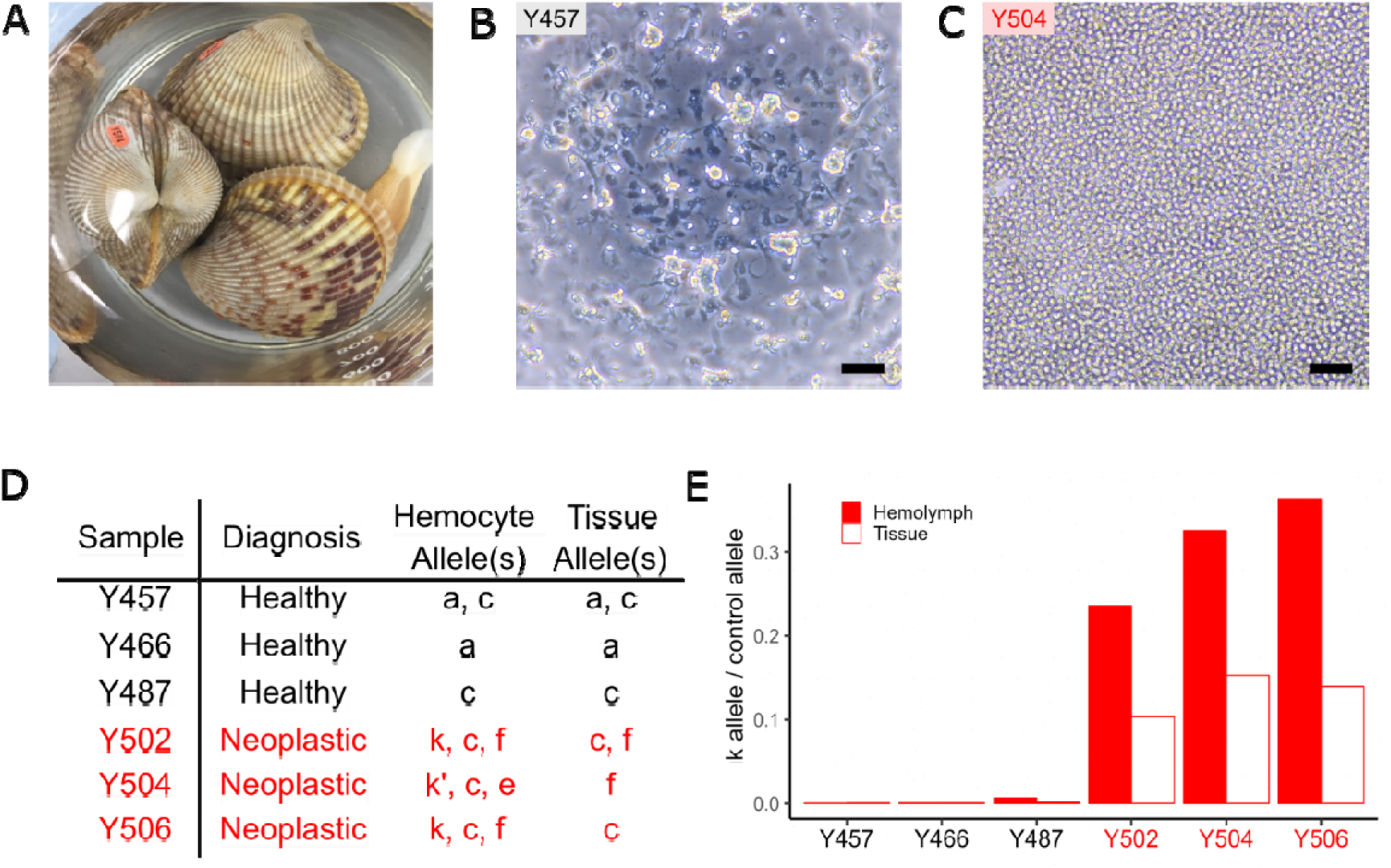
Identification of *Clinocardium nuttallii* bivalve transmissible neoplasia (CnuBTN1) as a transmissible cancer. A representative image of three basket cockles (*Clinocardium nuttallii*) is shown (*A*), collected from Agate Pass, Washington, USA in March 2019, including one animal with neoplasia (Y504). Phase-contrast microscopy of cells from hemolymph shows the difference in morphology between representative healthy (*B*) and neoplastic cockles (*C*). Scale bars are 50 µm. A fragment of the nuclear *α-tubulin* was amplified and sequenced from three healthy cockles (black) and three cockles identified as neoplastic through morphological examination of cells in hemolymph (red), and the alleles identified in the hemolymph and solid tissue of each cockle are shown (*D*). This shows the k and c allele in all three cases of cancer (DNA distance matrix of these alleles is found in Table S5). A qPCR assay was developed to determine the fraction of the total *α-tubuli n* alleles that were the cancer-associated k allele (*E*), showing a high fraction of the k allele in all three diseased animals, and a higher fraction of the k allele in the hemolymph than in the solid tissues, as expected for a BTN.

We extracted DNA from cells in the hemolymph taken from three cockles with DN (based on hemocyte morphology) and three “healthy” control animals (in which DN was not observed). We also sampled mantle tissue from each animal to compare genotypes. In transmissible cancers, the genotypes of the cancer cells do not match genotypes of the hosts, and instead they match those of other cancer cells from other host animals (Metzger & Goff, 2016; Metzger et al., 2015; Metzger et al., 2016). To test whether disseminated neoplasia in basket cockles is a BTN, we generated a set of intron-spanning primers in a conserved gene (*α-tubulin*) that would amplify a polymorphic region in all healthy and neoplastic samples. In the three healthy animals, one or two alleles were identified and the same one or two alleles were identified in DNA from both hemolymph and solid tissue samples within each healthy animal (alleles are given arbitrary letter names with identical alleles sharing the same letter, Figure 1D). In contrast, in all three cockles identified as neoplastic based on cell morphology, alleles found in the DNA extracted from hemolymph (mostly cancer) and from the solid tissue (mostly host) were different. Nearly identical alleles (alleles c and k/k’) were found in the neoplastic hemolymph samples for each of the three cockles. The k’ and k alleles differ by one unique single nucleotide variant (SNV) not observed in any healthy cockles, so it is likely due to somatic mutation within the cancer. We subsequently refer to these both as the k allele. These data confirm that disseminated neoplasia in *C. nuttallii* is a bivalve transmissible neoplasia (termed CnuBTN1).

Unexpectedly, we observed three alleles at this locus that were specific for the cancers instead of just two (Figure 1D). This could be due to a duplicated copy of the gene in the genome of the founder cockle that had the original cancer, or it could be due to horizontal transfer of nuclear DNA into the cancer lineage from host cells or another cancer lineage. One of the neoplastic samples (Y504) has a different third allele from the other two, (allele e instead of f). It is possible that this arose from a k allele that acquired a somatic mutation, but as the e allele has been observed in the germline of many normal cockles, it is possible that horizontal transfer of nuclear DNA may have occurred in these lineages, leading to this e allele. Based on the quantitation of the allele frequencies in the cancer samples, the three alleles appear to occur in roughly 1:1:1 proportions (Figure S1). This suggests that the detection of the three alleles is not due to a minor contamination and that they likely reflect the true genotype of the cancer cells at a triploid locus. Despite the variable third allele, the k and c alleles are present in all three cockle cancer samples, strongly suggesting that the neoplastic cells originated from a single cell lineage.

In order to quantify the amount of BTN-specific DNA in hemolymph and solid tissue samples and to generate an assay that can be used as a sensitive survey for the presence of BTN, we designed a qPCR primer pair specific for the cancer-associated *α-tubulin* k allele, as well as a control “universal” qPCR primer pair that amplifies the *α-tubulin locus* in all cockles. qPCR data for CnuBTN1-positive animals show that the putative cancer-associated k allele can be found in the solid tissue DNA samples of the neoplastic individuals, likely due to dissemination of BTN into host tissues; however, it is found in a much higher fraction in the DNA from the hemolymph samples, as would be expected for BTN (Figure 1E, (Metzger et al., 2015; Metzger et al., 2016)).

### Survey of Washington State cockles reveals evidence of two lineages of CnuBTN

In order to determine the extent of the spread of infectious cancer (CnuBTN) in basket cockle populations, we analyzed samples from a large population survey from Puget Sound and across Washington State (Dimond et al., 2022). Approximately 30 individuals were sampled from each of 11 different locations (n = 335). We used both slide-smear hemocytology (Vandepas et al., 2024) to analyze cell morphology and the cancer-specific qPCR assays to screen DNA from the hemolymph and tissue of each cockle. The slide-smear hemocytology method identified three cockles with clear evidence of advanced DN in this survey (SM26 and SM6 from Semiahmoo and PC2 from Penn Cove, Figure 2A-D).

**Figure 2.**
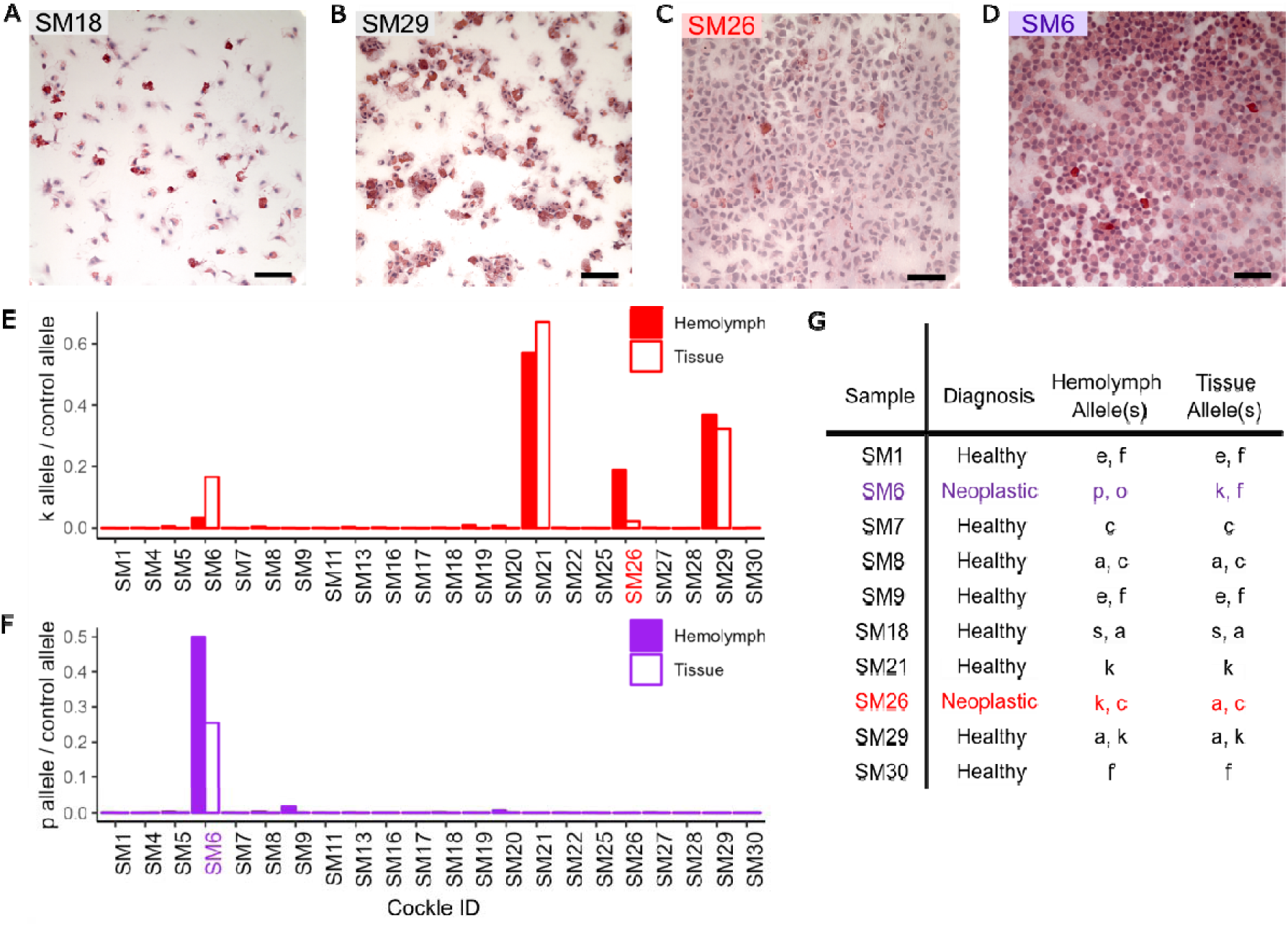
Screening of cockles from Semiahmoo, Washington reveals a second lineage o f CnuBTN. Cockles from Semiahmoo, Washington, USA, were analyzed by slide-smear hemocytology. Representative images from cockles diagnosed as healthy (*A*, *B*) and neoplastic (*C*, *D*) are shown (scale bar 50 µm). qPCR screening with the *α-tubulin* k-specific primers (*E*) showed that one neoplastic cockle was positive for the k allele in the hemolymph (SM26). The high level of k in both hemolymph and tissue in animals with normal hemocyte morphology (SM21 and SM29) show that these animals have the k allele in their germline. One cockle (SM6) was neoplastic by morphology, but negative for the k allele. The *α-tubulin* region was sequenced, and a p allele was identified that was specific for the cancer in this cockle (*F*). The region was cloned, and all alleles were sequenced for cancer samples and a subset of healthy cockles; all alleles of the cockles that were sequenced in this population are shown (*G*), with colors denoting healthy cockles (black), cockles with CnuBTN1 (red), and CnuBTN2 (purple).

From the collection at Semiahmoo, qPCR analysis of one of the cockles with DN (SM26) showed a high frequency of the putative cancer-associated k allele in the hemolymph and a lower frequency in the host tissues, similar to what was observed in DN+ individuals from Agate Pass (Figure 2C,E). Two cockles (SM21 and SM29) with normal hemocyte morphology were found to have a high frequency of the k allele in both hemolymph and tissue DNA (Figure 2B,E). This showed that the k allele was the endogenous germline allele in those animals, and that while the k allele can be used as a sensitive screen for CnuBTN, it is not specific (See Figure S3). As such, BTN can only be identified as likely CnuBTN1 if the hemolymph:tissue ratio of the k allele is higher than 1. Any samples meeting this screening criteria were verified through further sequencing.

Although morphological analysis of cells from cockle SM6 showed clear evidence of disseminated neoplasia (Figure 2D), the k allele was not present at a high fraction in the hemolymph. Amplification and sequencing of this cockle revealed that the cancer-specific alleles in SM6 were different from the host, but that, instead of alleles k and c, they were different alleles (p and o). Notably, the p allele is far more closely related to alleles found in other healthy individuals than it is to any allele found in the first lineage of CnuBTN. This shows that SM6 carried a different BTN lineage (CnuBTN2) that is independent of the first BTN lineage (now termed CnuBTN1) that was found in SM26 and the pilot samples from Agate Pass. A p allele-specific qPCR primer pair confirmed that this allele was found predominantly in the hemolymph of SM6, with a much lower level in the solid tissue of that individual, as expected for a BTN, but it was not confirmed in any other cockles (Figure 2F,G).

Cockles from the other 10 sites were also surveyed using qPCR to identify individuals with the k and p alleles indicative of CnuBTN1 and CnuBTN2, respectively, complementing the morphological analysis (Figure 3). Individuals identified as likely k- or p-allele positive through qPCR screening were further verified by cloning and sequencing of the α-*tubulin* locus. Overall, we confirmed the presence of CnuBTN1 in three locations in Puget Sound (Semiahmoo, Agate Pass, and Penn Cove), while CnuBTN2 was only found in Semiahmoo. Notably, the CnuBTN1 sample found in Penn Cove (PC2) matched the original samples from Agate Pass, in that it had three *α-tubulin* alleles (k, c, and f), while the sample from Semiahmoo (SM26) had only two observable alleles (k and c), and they were in a 1:2 ratio, suggesting that it was still a triploid locus, but with 2 copies of allele c and one of k, due to loss of heterozygosity, or some other mechanism (Figure S1).

**Figure 3.**
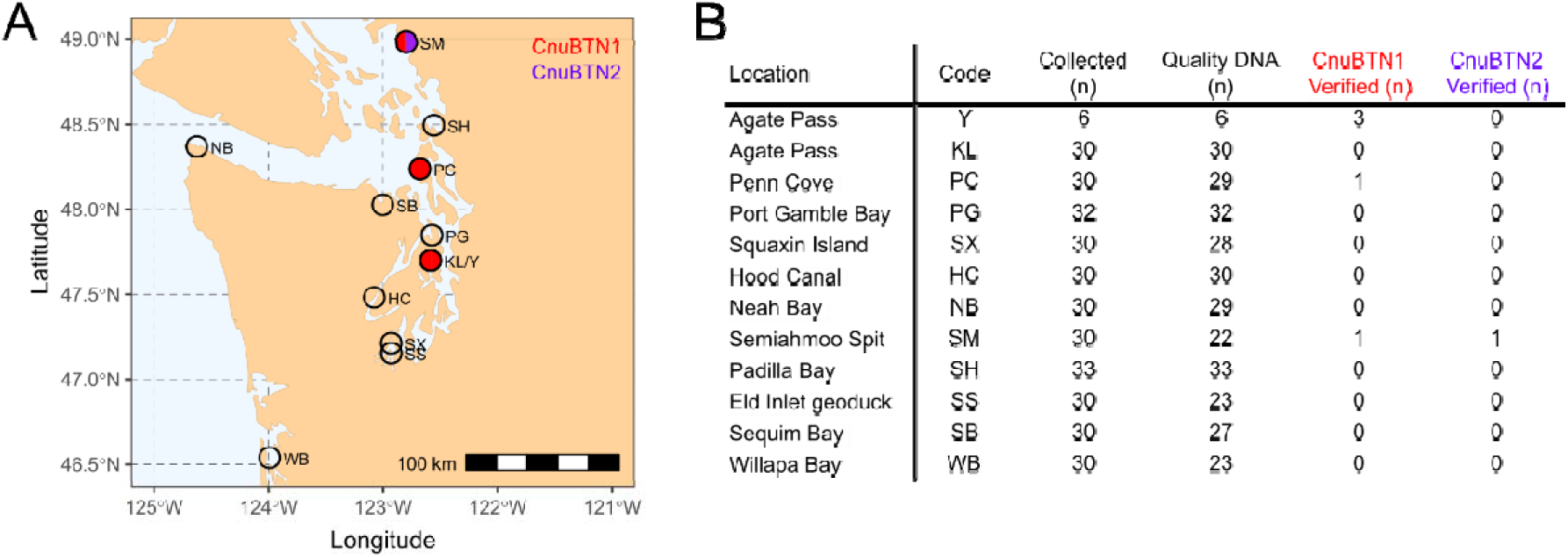
Screening of 11 sites in Washington State for BTN shows widespread and patch y distribution. Cockles collected from 11 different locations in Washington State were diagnosed for th presence of DN through cytology, and for two different lineages of transmissible cancer through qPCR and sequence analysis. (*A*) Map was made with R package rnaturalearthhire (https://github.com/ropensci/rnaturalearthhires). Each site is marked as having no BTN (clear), CnuBTN1 (red), CnuBTN2 (purple), or more than one (multiple colors). (*B*) The table lists the number of cockles collected, the number with extracted DNA of high enough quality to analyze, and the number of cases of each lineage of CnuBTN that were both identified through qPCR and were verified by sequencing of the *α-tubulin* locus.

Phylogenetic analysis of the α*-tubulin* alleles provides evidence supporting two independent CnuBTN lineages (Figure 4). To confirm this finding, we used 8 microsatellite primer pairs designed to identify polymorphic loci in basket cockles (Figure 5). Microsatellite alleles reveal differences between the individual samples of CnuBTN1, but all CnuBTN1 microsatellite genotypes cluster together, consistent with the hypothesis that all samples identified as CnuBTN1 come from a single BTN lineage. Additionally, the SM6 cancer microsatellite genotype is distinct from the CnuBTN1 cluster and from its host genotype, providing further evidence that this sample represents an additional independent lineage of BTN (CnuBTN2).

**Figure 4.**
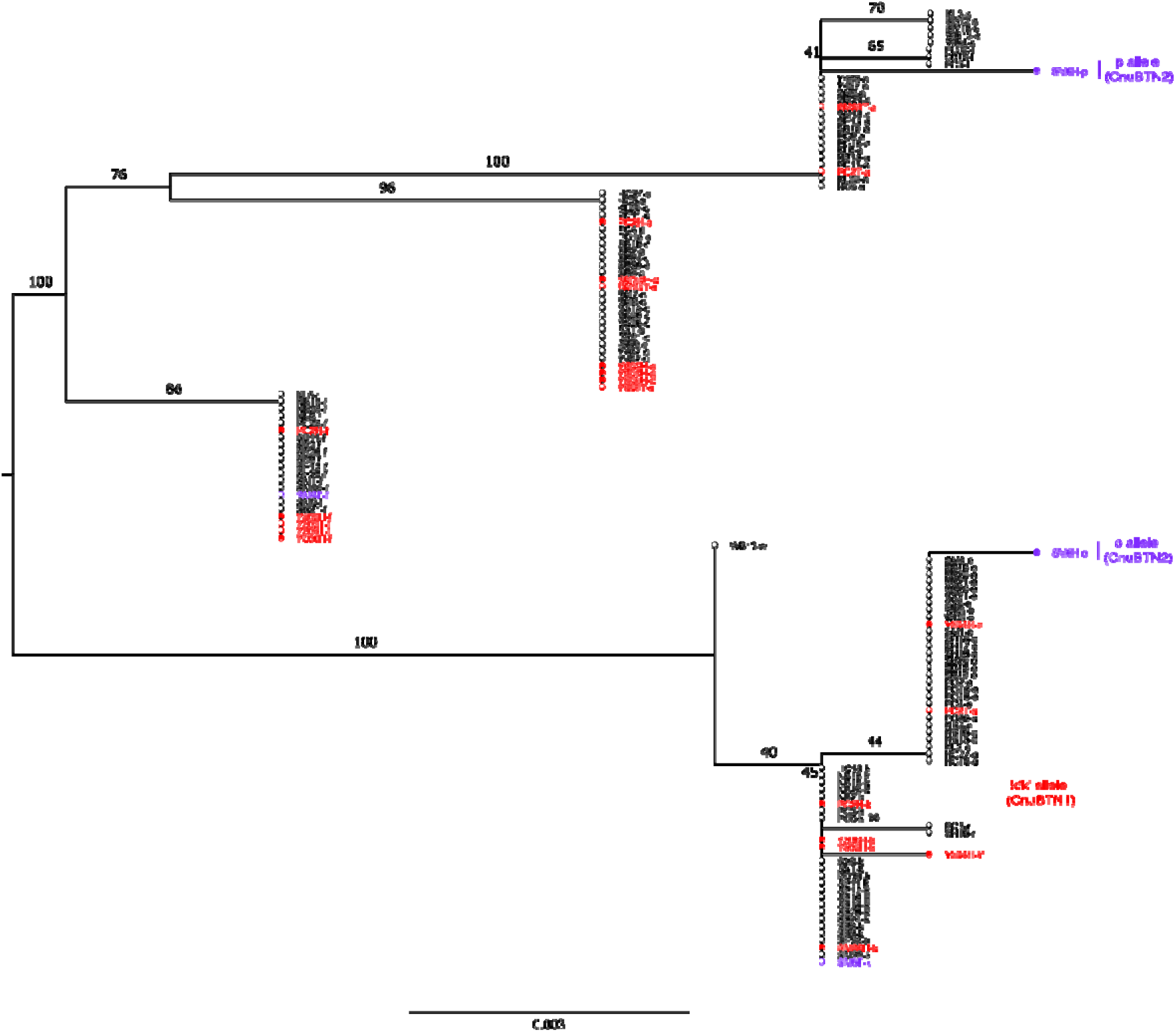
Phylogenetic analysis of *α-tubulin* sequence of DNA from BTN, host tissue, and healthy cockles. An intron-spanning region in the *α-tubulin* region was amplified and sequenced for all cockles that were identified through qPCR screening as likely BTN and for representative healthy animals from each collection. A phylogenetic tree was made with PhyML (Guindon et al., 2010; Lefort et al., 2017), rooted at the midpoint, with 100 bootstraps (values <30 not shown), using the GTR model (selected based on AIC). Alleles are marked with cockle ID and the allele name, and healthy cockles have 1-2 alleles (black open circles). Alleles from the host tissue of neoplastic cockles (colored open circles) are marked with “T” and alleles from cancer in hemolymph (colored filled circles) are marked with “H” (CnuBTN1, red; CnuBTN2, purple). Scale bar shows phylogenetic distance.

**Figure 5.**
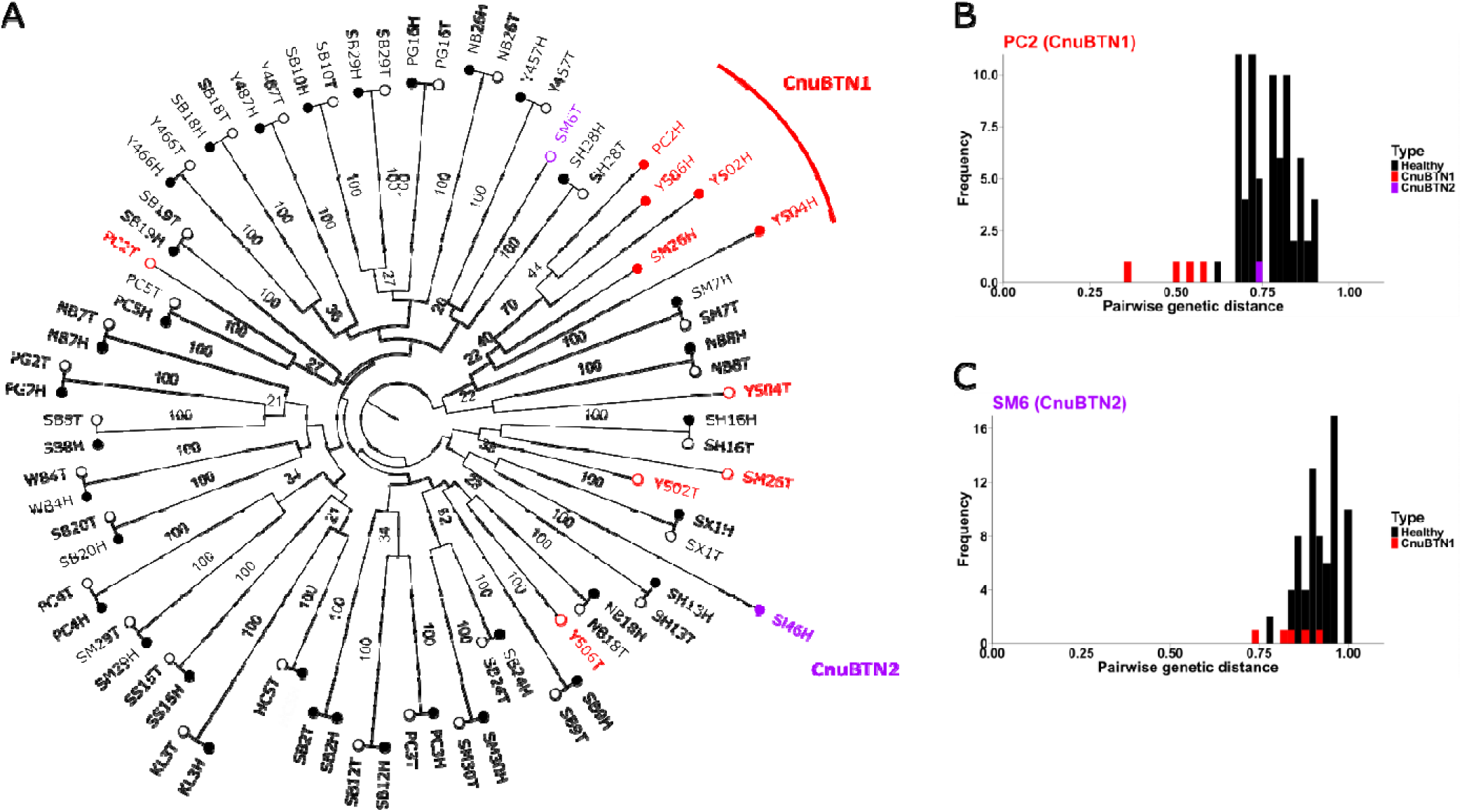
Microsatellite analysis of DNA from BTN, host tissue, and healthy cockles. Eight microsatellite loci were sequenced, and the allele sizes were identified from all samples with BTN and representative healthy animals from each collection (examples of microsatellit allele size calling are in Figure S2). (*A*) A phylogenetic tree was made from the allele sizes, using a binary marker model, rooted at the midpoint, with 100 bootstraps (values <20 not shown). Genotypes are marked with cockle ID and with “H” (hemocyte genotype, filled circles) or “T” (tissue genotype, open circles). Both BTN lineages cluster separately (Healthy cockles, black; CnuBTN1, red; CnuBTN2, purple). No scale bar is shown as the branch lengths using this binary model are normalized to the number of observed alleles and not to the number of loci, so they show relationships, but not an absolute genetic distance. Additionally, pairwise genetic distances were calculated from microsatellite data of all BTN, host, and healthy samples, using the co-dominant model. (*B*) A histogram is shown with pairwise distances between all samples and a representative sample of CnuBTN1 and (*C*) the sample of CnuBTN2. These data confirm that CnuBTN1 samples are more closely related to each other than to the other cancer lineage or to other healthy animals. Histograms of pairwise distances from all CnuBTN1 samples are in Figure S4.

### Horizontal transfer of mitochondrial genomes in CnuBTN1

Our analyses of multiple nuclear markers show that CnuBTN1 samples form a clonal lineage; however, sequencing of mitochondrial DNA revealed that every sample of BTN observed in this study had a different mitogenome sequence. Sequences of both *cytochrome c oxidase* (*mtCOI*) and *cytochrome b* (*mtCYTB*) show that the BTN-associated mitochondrial alleles are different from host alleles, but the CnuBTN1 sequences do not support a single origin for the mitochondria of this cancer lineage (Figure 6). Data from both mitochondrial loci support the HT1 event to Y502, and considering the data from both loci, a minimum of two separate horizontal transfer events within CnuBTN1 would be needed to explain the mitochondrial sequence divergence (HT1 for cockle Y502 and HT2 for cockle Y504). Additional complete mitogenome sequencing of cancer and healthy samples will be required to fully resolve these HT events and to determine whether the differences observed in other samples (such as PC2 and Y506) are due to somatic mutations from a common original mitogenome or due to additional HT events. Additionally, at least three separate mitogenome alleles were observed in the case of Y504, suggesting either simultaneous infection with multiple divergent clones, or ongoing heteroplasmy and recombination.

**Figure 6.**
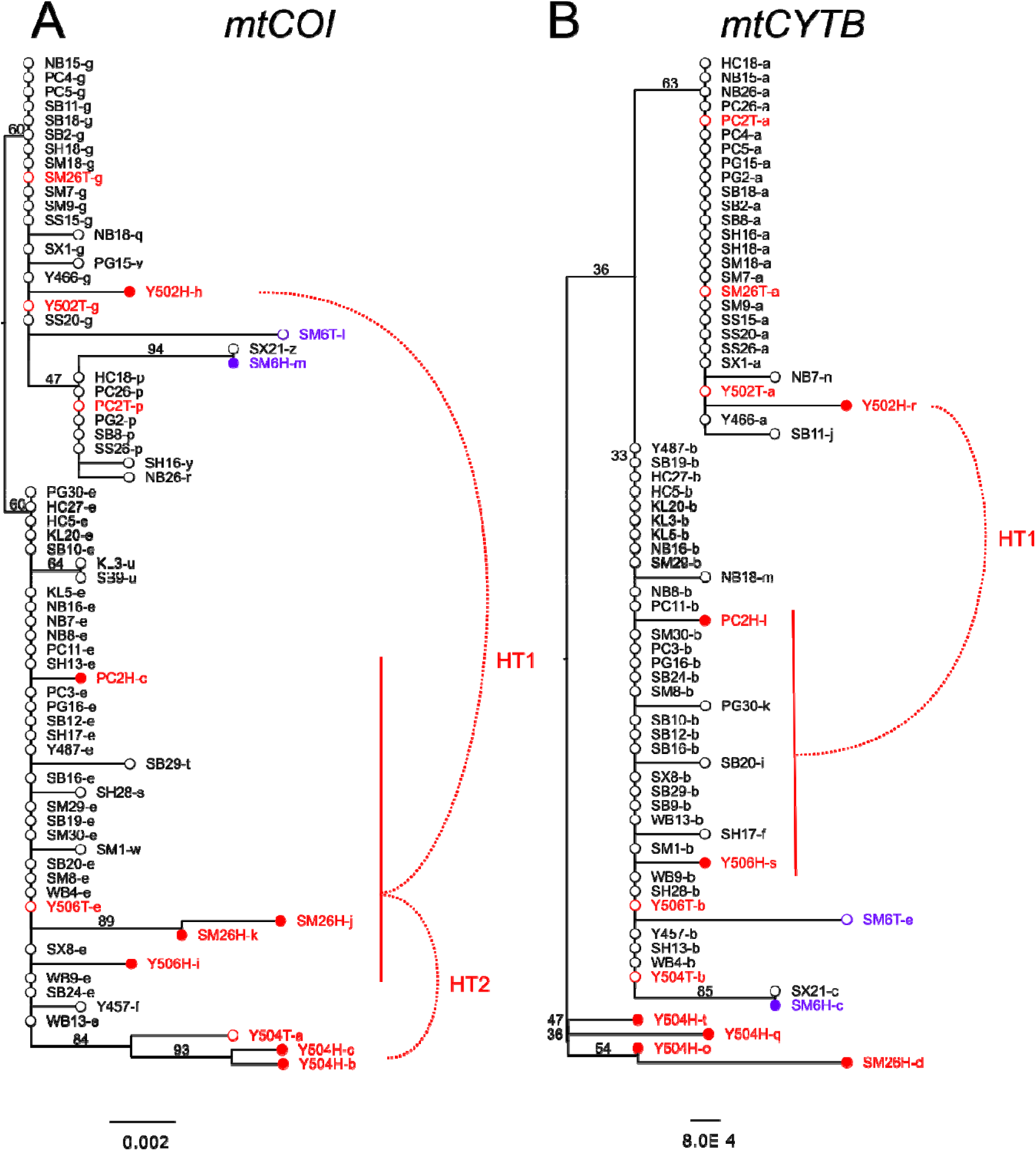
Phylogenetic analysis of mitochondrial loci (*mtCOI* and *mtCYTB*) Two loci from the mitochondrial genome, *mtCOI* (*A*) and *mtCYTB* (*B*) were amplified and sequenced for each animal with BTN and for representative healthy samples from each collection (healthy, black; CnuBTN1, red; CnuBTN2, purple). A phylogenetic tree was made for each locus using PhyML (Guindon et al., 2010; Lefort et al., 2017), rooted at the midpoint, with 100 bootstraps (values <30 not shown) using the TN93+G model (selected as it was one of the top 3 models based on AIC for both loci). The mtDNA sequences of both loci from all CnuBTN1 samples (red, filled circles) are different from each other, and more closely related to those sequences found in healthy cockles, suggesting multiple horizontal transfer (HT) events. Some different sequences could be due to HT or due to divergence from a common cancer allele (ie. the differences between cancer sequences in PC2, Y506, and SM26 could be due to the cancer originating with *mtCOI* -e & *mtCYTB* -b alleles), but for at least two cases, horizontal transfer from a different healthy mitogenome is the most likely explanation for the sequence divergence (HT1 to Y502 and HT2 to Y504). Multiple cancer-associated alleles were found in SM26 and Y504. All cancer-associated alleles are shown on the tree and may represent multiple related cancers in the same cockle or a single cancer lineage with heteroplasmy. Scale bars show phylogenetic distance.

## DISCUSSION

This study provides the first report of transmissible cancer in basket cockles (*C. nuttallii*), with evidence from multiple nuclear loci (α*-tubulin* alleles and 8 microsatellite loci) showing two independent lineages of *C. nuttallii* BTN (CnuBTN1 & 2). In conventional cancers, somatic mutations do occur, but there is no method to explain the simultaneous acquisition of the same variants in a housekeeping gene, or to explain the convergence of multiple microsatellite loci to a specific size pattern (some becoming larger and some smaller than the host alleles). However, this is what would be expected in a transmissible cancer, as the cancer cells from a single lineage that are found infecting different animals all arose from a single individual animal. This strategy of finding genetic markers in a cancer (not connected to any specific driver mutations) that are different from the hosts and identical across multiple individuals is the primary method used to identify transmissible cancers in dogs, devils and other bivalve species (Garcia-Souto et al., 2022; Metzger et al., 2015; Metzger et al., 2016; Michnowska et al., 2022; Murgia et al., 2006; Pearse & Swift, 2006; Pye et al., 2016; Rebbeck et al., 2009; Yonemitsu et al., 2019).

One limitation of this study is the observation that the CnuBTN1 qPCR diagnostic assay, first thought to be cancer-specific, amplifies an allele later found in some healthy animals at several locations in Puget Sound. These cancers originally arise from normal cockles, so this is to be expected, but it highlights the difficulty in relying on a single genetic marker for disease diagnosis. Recently, a similar study found that an *EF1-* α allele previously only found in the mussel *Mytilus trossulus* BTN (MtrBTN1) is also present in the germline of non-neoplastic animals in Northern Europe (Skazina et al., 2021; Skazina et al., 2023). This suggests that the cancer may have arisen from the Northern Europe mussel population, but also renders that marker non-specific in animals with that allele. In the absence of better markers and more complete surveys of healthy genetic diversity, researchers must use a combination of diagnostic assays, or use the cancer:host ratio to determine whether a sample is host to a transmissible cancer lineage, as we have done in this study.

Additionally, our finding of multiple lineages of CnuBTN relied on the identification of neoplastic cells through morphological analysis, demonstrating that a diagnostic approach based on genetic markers alone can miss the presence of additional lineages of transmissible cancer in bivalve populations.

Unexpectedly, while both sequence data from the *α-tubulin* locus and microsatellite allele sizes from multiple loci strongly support a single clonal lineage of CnuBTN1, mitochondrial DNA sequences are not consistent with a clonal mitogenome origin. Horizontal transfer of mitochondrial DNA has been reported in the canine transmissible venereal tumor of dogs (Rebbeck et al., 2011; Strakova et al., 2020), as well as more recently in the BTNs of *Mytilus trossulus* (Yonemitsu et al., 2019) and the common cockle, *Cerastoderma edule*(Bruzos et al., 2023). Our observations of multiple horizontal transfer events within just a few samples of the CnuBTN1 lineage suggest that cockles may have a predisposition to mitochondrial transfer. For example, the MtrBTN2 lineage has now been observed in four different species in Europe, South America, and Asia, and, to date, only one horizontal transfer event has been documented. No horizontal transfers have yet been observed in *Mya arenaria* BTN (Hart et al., 2023). It is unclear in these cases whether mitogenome replacement is due to selection for more functional mitochondria after degradation of the mitogenome through the Muller’s Ratchet of somatic mutation or due to a selfish benefit to the mitochondrial genome itself (as suggested for CTVT based on the repeated transfer of alleles with the same mutation in the mitochondrial control region (Strakova et al., 2020)). The horizontal transmission of DNA is not limited to transmissible cancers. It has been suggested that this sort of parasexual recombination may be important for the evolution of human cancers (Miroshnychenko et al., 2021), and human cancer cells have been shown to take up functional mitochondria from cancer-associated host cells in multiple studies (Griessinger et al., 2017; Liu et al., 2021). Based on our observation of frequent replacement of mitogenomes in cockles, these lineages are critical models for understanding the process of transfer and the selection pressures involved in cancer mitochondrial evolution.

Bivalve transmissible cancer is often a fatal disease, and in some cases has been reported to cause severe population loss of more than 90% (Farley et al., 1991; Muttray et al., 2012). It is clear that multiple lineages of CnuBTN are present in the basket cockle populations in Puget Sound, but it is not yet known whether they are causing mortality events or how they are being spread from one location to another. Further research on this newly described infectious agent will provide an important current baseline of geographic and seasonal occurrences of this disease, enable understanding of how it spreads through the environment, how future climate conditions may affect incidence and severity, how it affects the survival and selective pressures on cockles, and how aquaculture and ecological management strategies might mitigate the disease in the future.

## Acknowledgements

We are grateful for the assistance of many people who collected cockles or helped facilitate collections, including Viviane Barry, Jodie Toft, Adrianne Akmajian, Julie Barber, Kevin Cagey, Tamara Gage, Jim Gibbons, Donovan Henry, Neil Harrington, Paul Harris, Megan Hintz, Nik Matsumoto, James McArdle, Matt Nelson, Blair Paul, Andy Pavone, Ralph Riccio, Marilyn Sheldon, Eric Sparkman, and Liz Tobin. We are thankful for the laboratory assistance in preparing and processing tissue samples by Paul Blodgett, Josh Bouma, Lisa Crosson, Claire Everett, Mackenzie Gavery, and Crystal Simchick, and support from Frederick Goetz. We are grateful to the Benaroya Research Institute’s Histology Core for technical support. Many specimen collections were made on Tribal reservations and/or usual and accustomed fishing grounds, and we are grateful to the Jamestown S’Klallam Tribe, the Lummi Nation, the Makah Nation, the Port Gamble S’Klallam Tribe, the Skokomish Tribe, the Suquamish Tribe, the Squaxin Island Tribe, and the Swinomish Indian Tribal Community for granting permission to collect, and in most cases performing the collections. Gratitude to Suquamish Tribal Members, Azure Boure and Luther Mills, Jr, who provided indigenous historical and contemporary frameworks for locating, harvesting and community subsistence, of local cockle populations. We also thank the Seattle Shellfish Company for collection of cockles from Eld Inlet geoduck tubes. Additional collections were made under Washington Department of Fish and Wildlife Scientific Collection Permit 19-319.

## Conflict of Interest Statement

The authors declare no conflicts of interest. Data Accessibility and Benefit-Sharing:

All data are available as supplementary files. All sequences are available on NCBI Genbank (BioProject PRJNA1082776). Raw files with fragment analysis data from microsatellites can be found at https://www.bco-dmo.org/project/936145 (doi:10.26008/1912/bco-dmo.940678.1).

---

**Figure S1.**
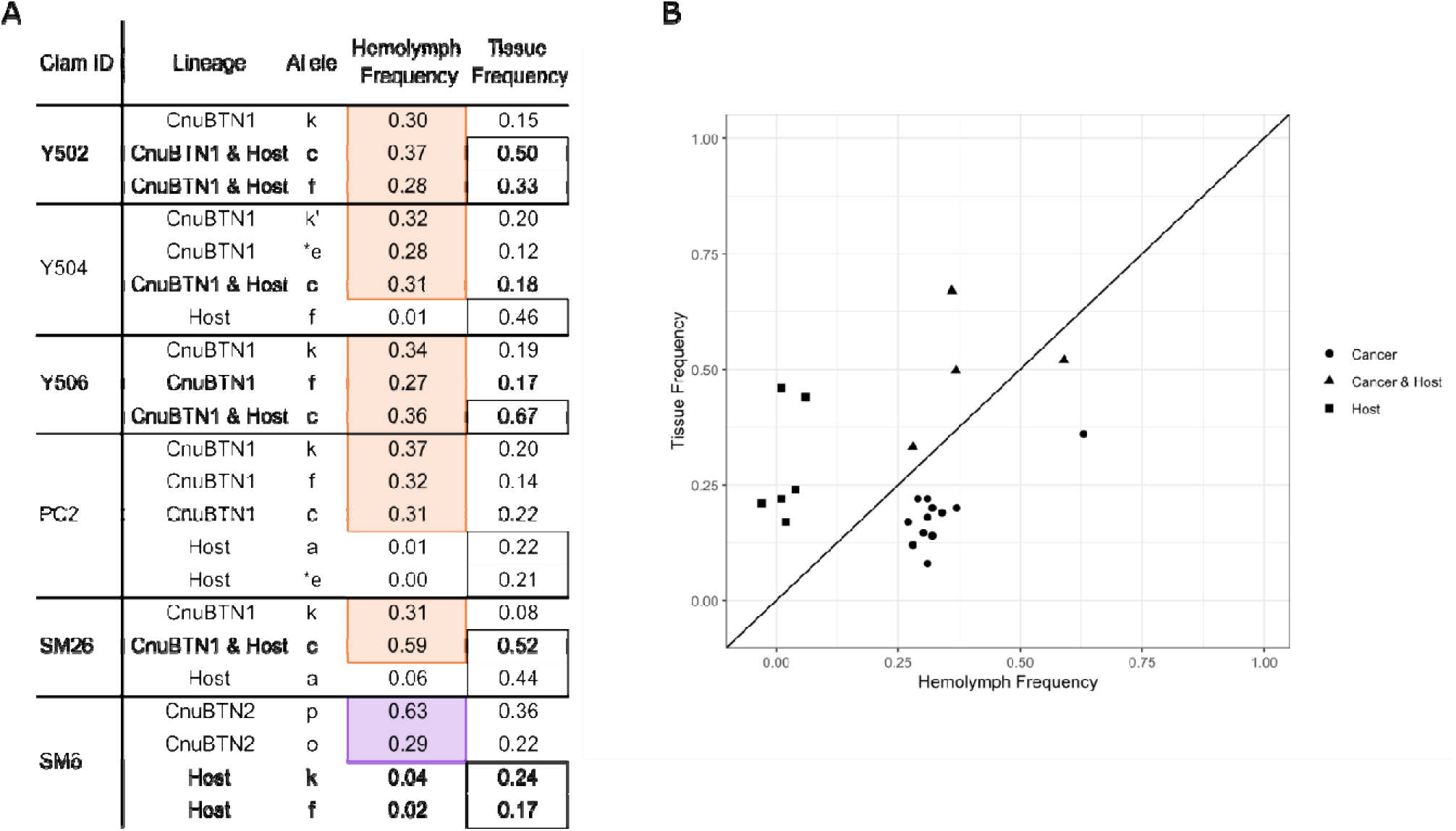
Quantification of allele frequencies of *α-tubulin*. The PCR products from the intron-spanning region of *α-tubulin* amplified from the hemolymph DNA and solid tissue DNA of cockles with BTN were sequenced using nanopore technology and the frequency of each allele was calculated based on the average of the frequency of detection of SNVs that are unique to each allele (*A*). The alleles that were identified a originating from the host are marked with an open black box, and the alleles that were identified as originating from the cancer are shaded with color (CnuBTN1, red; CnuBTN2, purple). As the frequency of each allele was measured by a different set of SNVs the sum of the alleles frequencies in each sample is not exactly 1 (range = 0.93-1.05). Allele e (marked with an *) lacked any unique SNVs, but it was different from the k allele by a single SNV. In this case, the e allele frequency for each sample was calculated by taking the average allele frequency of SNVs common to both k and e in those samples and subtracting the frequency of the SNV found only in k and not e. For the PC2 host samples, this calculation yields a value of -0.03, so it was corrected to 0.00 for the table, as no real negative frequencies are possible and this just represents a minor artifact due to subtracting two similar values with experimental error. A scatterplot of the values of allele frequency in tissue and hemolymph DNA are shown (B) to illustrate which values were called as originating from the host (squares), from the cancer (circle) or present in both host and cancer (triangle). A line of slope = 1 is shown for reference.

**Figure S2.**
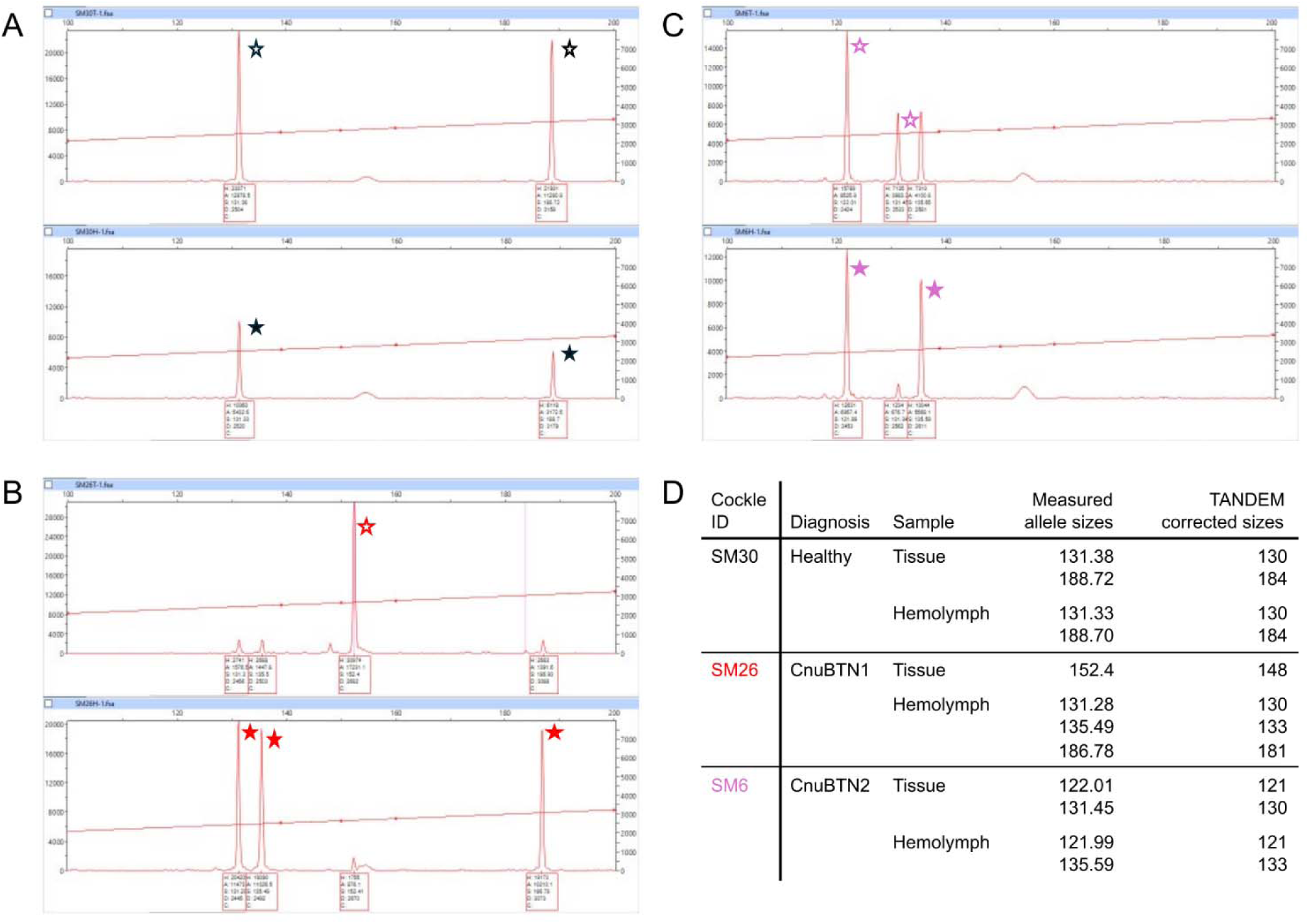
Examples of calling alleles from raw microsatellite allele data in healthy and neoplastic clams. For each microsatellite locus analyzed, the fragment analysis files (.fsa) were inspected using the Peak Scanner Software. A representative locus is shown here (Cnu58), and the raw values for the solid tissue DNA (top) and hemolymph DNA (bottom) are shown for one representative healthy cockle (*A*, SM30), one representative cockle with CnuBTN1 (*B*, SM26), and the cockle with CnuBTN2 (*C*, SM6). Stars mark the alleles (open for the solid tissue genotype and filled for hemolymph genotype). For all healthy cockles, these alleles match, as shown in (*A*), but for cockles with BTN there are different alleles observed representing the host and cancer genotypes. The exact measured values were corrected to integer values using TANDEM, and the results are shown in the table (*D*).

**Figure S3.**
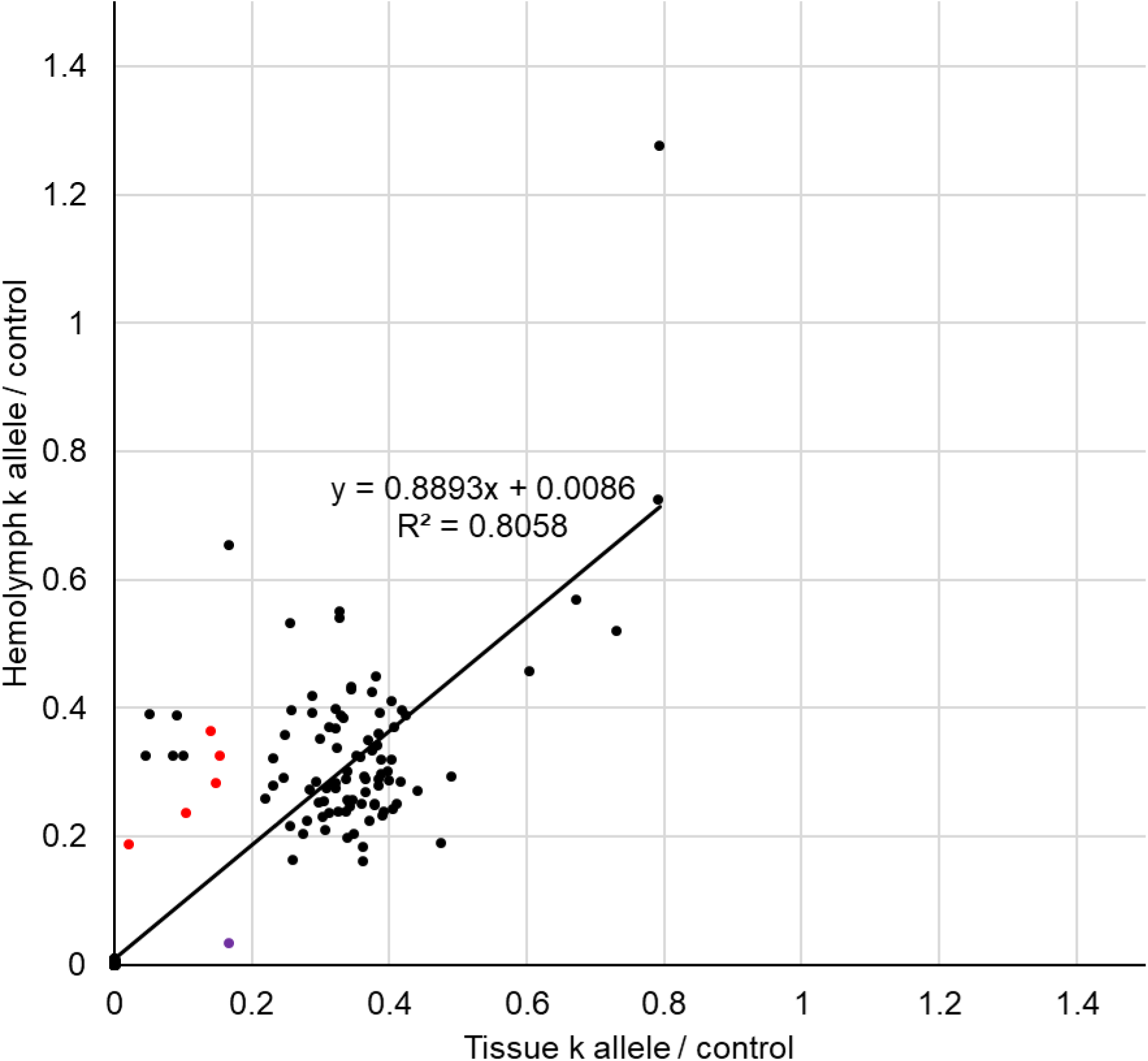
Comparison of the CnuBTN1-specific k allele in hemolymph and tissue DNA of all cockle samples. The *α-tubulin* k allele was identified as specific to the CnuBTN1 lineage in the pilot set of samples from Agate Pass, but in screening of other samples it was found that a large subset of healthy cockles has the k allele in their germline (See Figure 2). The ratio of the amplification of the k allele compared to amplification with the control primers is shown here for the hemolymph DNA and tissue DNA of all samples collected that passed quality control (n=311). The majority of samples do not have the k allele (211 have a ratio of <0.01 for both hemolymph and tissue). Samples with a higher amount of k allele in the hemolymph than in tissue were further analyzed as possible cases of CnuBTN1 (verified cases are shown in red). Most samples with the k allele have roughly equivalent levels of k allele in hemolymph and tissue, corresponding to healthy cockles with the k allele as either heterozygous or homozygous in their germline (black dots). The SM6 cockle with CnuBTN2 has the k allele in the host genotype but not the cancer (purple dot). Line shows linear regression of healthy samples.

**Figure S4.**
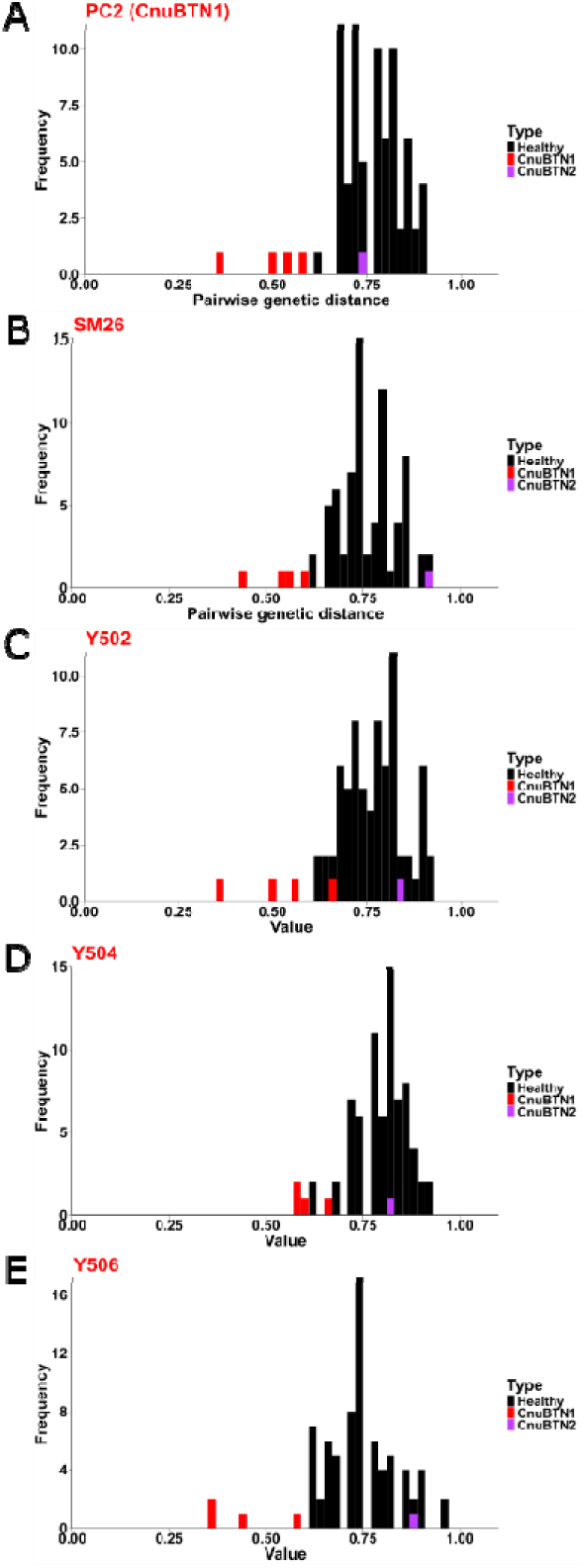
Pairwise microsatellite genetic distances from all CnuBTN1 samples to all other healthy samples and CnuBTN2. Eight microsatellite loci were sequenced, and the allele sizes were identified from all samples with BTN and representative healthy animals from each collection (examples of microsatellit allele size calling are in Figure S2). Pairwise genetic distances were calculated from microsatellite data of all BTN, host, and healthy samples, using the co-dominant model, as in Figure 5. Histograms are shown with pairwise distances between each sample of CnuBTN1 (*A*, PC2; *B*, SM26; *C*, Y502; *D*, Y504; and *E*, Y506) and all other samples (including normal healthy DNA samples and host tissue samples from cockles with BTN, black; and the CnuBTN2 cancer sample, purple). Note that there was evidence from the sequencing of the *α-tubulin* locu suggesting that there has been horizontal transfer of nuclear DNA in the cancer from sample Y504. The microsatellite analysis is consistent with this hypothesis, as Y504 is the most distantly related of all the samples of CnuBTN1.

**Table S1.** Locations and dates of cockle collection.

| **Location** | **Beach Name** | **Code** | **Collection Date** | **Tribe/Agency** | **GPS Coordinates** |
| --- | --- | --- | --- | --- | --- |
| Agate Pass, WA | Kiana Lodge | Y | 3/18/2019 | Suquamish | 47.699608, -122.581537 |
| Penn Cove, WA | Monroe Landing | PC | 10/28/2019 | Swinomish | 48.237857, -122.67545 |
| Agate Pass, WA | Kiana Lodge | KL | 10/29/2019 | Suquamish | 47.699608, -122.581537 |
| Port Gamble Bay, WA | Gravel Plot, PGST Rez | PG | 10/29/2019 | Port Gamble S'Klallam | 47.847462, -122.570677 |
| Willapa Bay, WA | Stony Point Growing Area | WB | 11/13/2019 |  | 46.680, -123.959 |
| Squaxin Island, WA | S21 | SX | 11/14/2019 | Squaxin | 47.213164, -122.926208 |
| Eld Inlet geoducks tubes, WA | Schmidt | SS | 11/14/2019 | Seattle Shellfish | 47.155632, -122.925787 |
| Hood Canal, WA | Eagle Creek | HC | 11/14/2019 | Skokomish | 47.48232, -123.0747 |
| Neah Bay, WA | Front Beach | NB | 12/11/2019 | Makah | 48.36940, -124.62356 |
| Sequim Bay, WA | Blyn | SB | 12/11/2019 | Jamestown S'Klallam | 48.027609, -123.000763 |
| Semiahmoo Spit, WA | Semiahmoo Spit | SM | 12/12/2019 | Lummi/ PSRF | 48.98332, -122.79347 |
| Padilla Bay, WA | March's Point/Ship Harbor | SH | 1/9/2020 | PSRF/ Swinomish | 48.4951, -122.5539 |

**Table S2.**
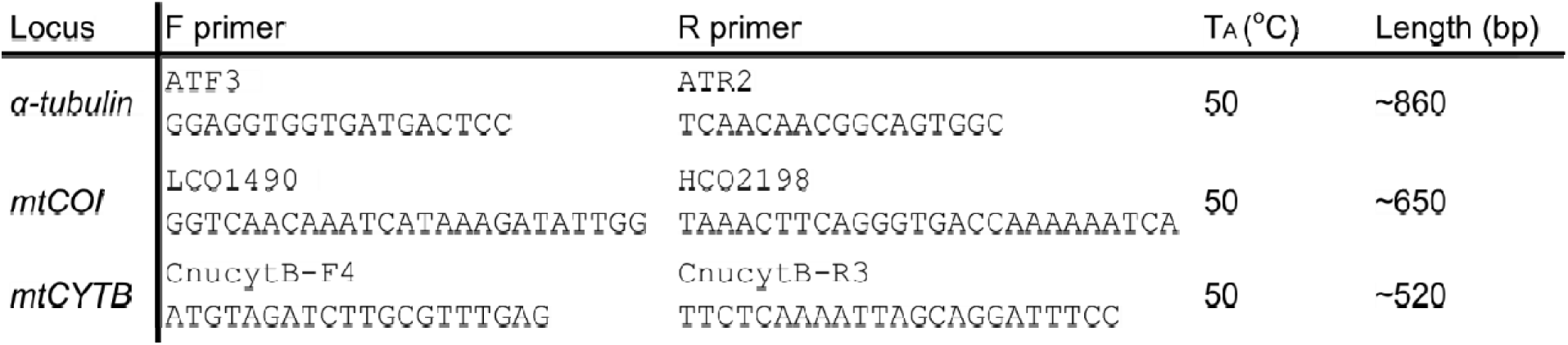
Primers used for amplification and sequencing.

**Table S3.**
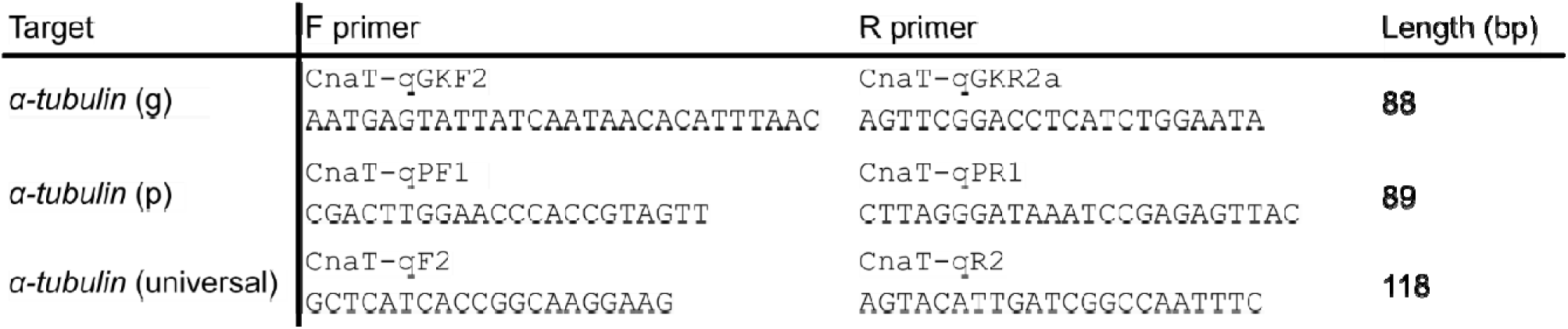
Primers used for allele-specific qPCR of *α-tubulin*.

**Table S4.**
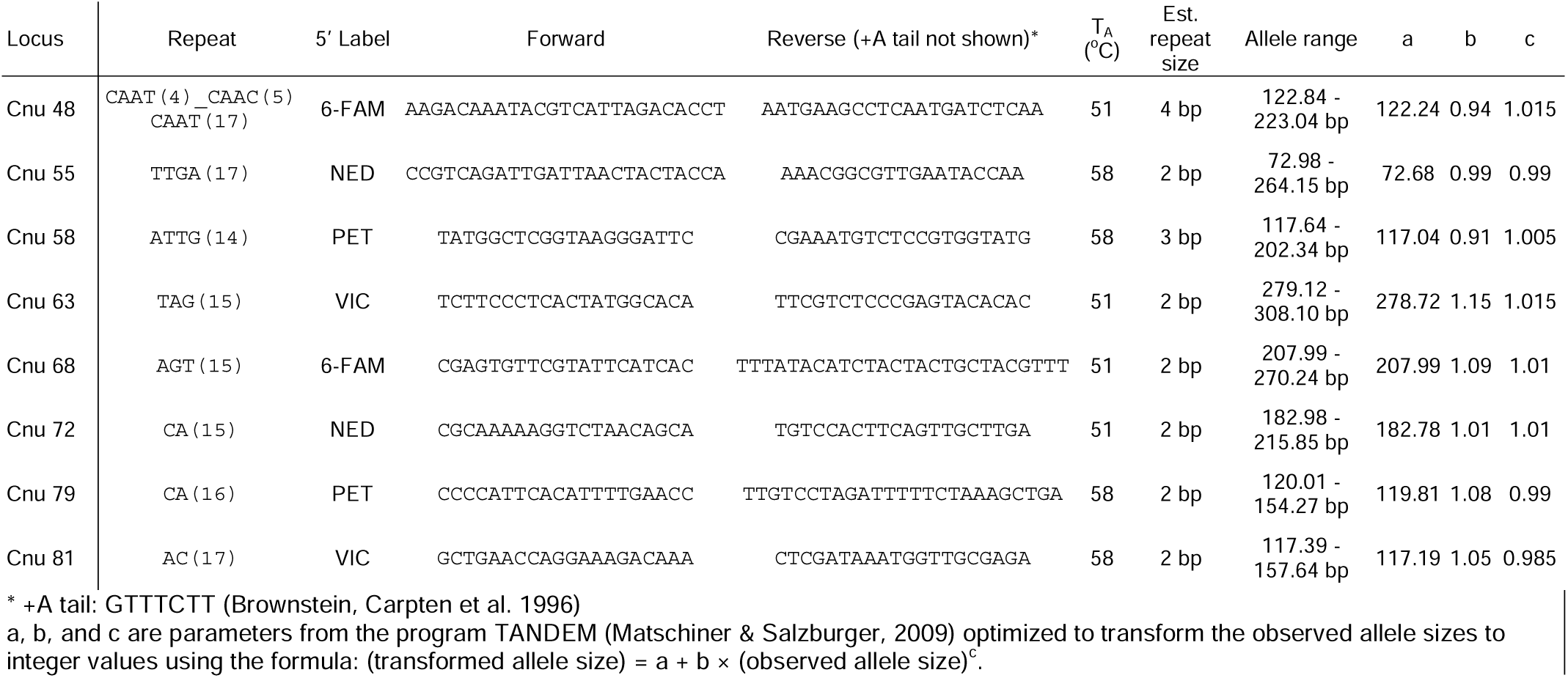
Primers for microsatellite amplification.

**Table S5.** DNA distance matrix for α-tubulin alleles sequenced from Agate Pass (collection Y)

|  | a | c | e | f | k | k' |
| --- | --- | --- | --- | --- | --- | --- |
| a | 0 | 0.0117 | 0.0164 | 0.0105 | 0.0176 | 0.0188 |
| c | 0.0117 | 0 | 0.014 | 0.0081 | 0.0152 | 0.0163 |
| e | 0.0164 | 0.014 | 0 | 0.0105 | 0.0012 | 0.0023 |
| f | 0.0105 | 0.0081 | 0.0105 | 0 | 0.0116 | 0.0128 |
| k | 0.0176 | 0.0152 | 0.0012 | 0.0116 | 0 | 0.0012 |
| k' | 0.0188 | 0.0163 | 0.0023 | 0.0128 | 0.0012 | 0 |

